## Supplementary Text for "Bayesian clustering with uncertain data"

### Bayesian clustering with uncertain data: Supplementary Information

#### 1 Gibbs equations

In this section we derive the conditional posterior distributions for each variable in the model, which are required for Gibbs sampling.

For convenience, we use vector notation to indicate all variables of a given type:

$$\begin{aligned}\boldsymbol{x} &= (x_1, \ldots, x_n)\\ \boldsymbol{z} &= (z_1, \ldots, z_n)\\ \boldsymbol{c} &= (c_1, \ldots, c_n)\\ \boldsymbol{\pi} &= (\pi_1, \ldots, \pi_K)\\ \boldsymbol{\mu} &= (\mu_1, \ldots, \mu_K)\\ \boldsymbol{\rho} &= (\rho_1, \ldots, \rho_K)\end{aligned}$$

We also introduce $\boldsymbol{c_{-i}}$ to represent all the indicators except the $i$th indicator i.e. $\boldsymbol{c_{-i}} = (c_1, \ldots, c_{i-1}, c_{i+1}, \ldots, c_n)$. We introduce $n_k$ as the number of points in cluster $k$ and $n_{-i,k}$ as the number of points in cluster $k$ excluding the $i$th point.

##### 1.1 Distribution of cluster membership indicators $c_i$

The conditional distribution of indicators $c_i$ is given by

$$p(c_i|\boldsymbol{c_{-i}}, \boldsymbol{x}, \boldsymbol{z}, \boldsymbol{\pi}, \boldsymbol{\mu}, \boldsymbol{\rho}, \alpha, K)$$

which, after marginalising out the cluster parameters $\boldsymbol{\mu}, \boldsymbol{\rho}$ and using $\theta_0 = (\mu_0, \kappa_0, \alpha_0, \beta_0)$ to group together the constant hyperparameters of the cluster parameters to ease notation is equivalent to:

$$p(c_i|\boldsymbol{c_{-i}}, \boldsymbol{x}, \boldsymbol{z}, \boldsymbol{\pi}, \alpha, K, \theta_0)$$

##### 1.1.1 Prior

Let us first consider the prior $p(\boldsymbol{c}|\boldsymbol{\pi}, \alpha, K)$. First, note that we can marginalise (Section 3.1) the cluster weights $\boldsymbol{\pi}$ to get:

$$p(\boldsymbol{c}|\alpha, K) = \frac{\Gamma(\alpha)}{\Gamma(n+\alpha)} \prod_{k=1}^{K} \frac{\Gamma\left(n_k + \alpha/K\right)}{\Gamma(\alpha/K)} \tag{1}$$

We can then (Section 3.2) use this formula to calculate the prior conditional on the other indicators:

$$\begin{aligned} p(c_i = k|\boldsymbol{c_{-i}}, \alpha, K) &= \frac{p(c_i = k, \boldsymbol{c_{-i}}|\alpha, K)}{\sum_{l=1}^{K} p(c_i = l, \boldsymbol{c_{-i}}|\alpha, K)} \\ &= \frac{n_{-i,k} + \frac{\alpha}{K}}{n - 1 + \alpha} \end{aligned} \tag{2}$$

**Prior in infinite limit** We now take the limit as the number of clusters $K$ tends to infinity. The formula above includes the term $n_{-i,k}$ and thus we need to consider two different cases. The simplest is where $n_{-i,k} > 0$ i.e. there is another point with the label $k$ already. For this case we simply take the limit as $K$ tends to infinity and get:

$$\text{if } n_{-i,k} > 0: \quad p(c_i = k|\boldsymbol{c_{-i}}, \alpha) = \frac{n_{-i,k}}{n - 1 + \alpha}$$

When $n_{-i,k} = 0$, i.e. there are no existing points in cluster $k$ we are creating a new cluster. It is convenient to combine all the subcases $p(c_i = k|\boldsymbol{c_{-i}, \alpha})$ for all clusters $k$ which are currently empty, i.e. all $k$ satisfying $\forall j \neq i, c_j \neq k$. When we combine all these subcases, this case simply becomes '*create a new cluster*'. By combining all these subcases and taking the limit we get the following:

$$\begin{aligned} p(\forall j \neq i, c_i \neq c_j|\boldsymbol{c_{-i}}, \alpha) &= \lim_{K\to\infty} \sum_{k=K_{\text{rep}+1}}^{K} \frac{n_{-i,k} + \frac{\alpha}{K}}{n - 1 + \alpha} \\ &= \lim_{K\to\infty} \sum_{k=K_{\text{rep}+1}}^{K} \frac{\frac{\alpha}{K}}{n - 1 + \alpha} \\ &= \lim_{K\to\infty} \frac{\left(K - K_{\text{rep}}\right)\frac{\alpha}{K}}{n - 1 + \alpha} \\ &= \frac{\alpha}{n - 1 + \alpha} \end{aligned}$$

where $K_{\text{rep}}$ is the number of non-empty clusters. The final line holds since as $K$ tends to infinity, $\frac{K - K_{\text{rep}}}{K} \to 1$ i.e. most clusters will be unoccupied since $n$ remains finite.

To summarise, conditional on all the other indicators, a point starts a new cluster of its own with weight $\alpha$ and joins each of the existing clusters $k$ with weight $n_{-i,k}$. Thus a point is more likely (a priori) to join a cluster with a large number of other points in it already.

##### 1.1.2 Posterior

We want to calculate the probability that we assign point $i$ to cluster $k$ given all the other variables. Using Bayes' Theorem, we can rewrite the desired probability as follows:

$$\begin{aligned} p(c_i = k|\boldsymbol{c_{-i}}, \alpha, \boldsymbol{z}, \boldsymbol{x}, \theta_0) &= \frac{p(c_i = k|\boldsymbol{c_{-i}}, \alpha)}{p(\boldsymbol{z}, \boldsymbol{x}|\boldsymbol{c_{-i}}, \alpha, \theta_0)} p(\boldsymbol{z}, \boldsymbol{x}|c_i = k, \boldsymbol{c_{-i}}, \alpha, \theta_0) \\ &= \frac{p(c_i = k|\boldsymbol{c_{-i}}, \alpha)}{p(\boldsymbol{z}, \boldsymbol{x}|\boldsymbol{c_{-i}}, \alpha, \theta_0)} p(\boldsymbol{x}|\boldsymbol{z}, c_i = k, \boldsymbol{c_{-i}}, \alpha, \theta_0) p(\boldsymbol{z}|c_i = k, \boldsymbol{c_{-i}}, \alpha, \theta_0) \\ &= \frac{p(c_i = k|\boldsymbol{c_{-i}}, \alpha)}{p(\boldsymbol{z}, \boldsymbol{x}|\boldsymbol{c_{-i}}, \alpha, \theta_0)} p(\boldsymbol{x}|\boldsymbol{z}) p(\boldsymbol{z}|c_i = k, \boldsymbol{c_{-i}}, \alpha, \theta_0) \end{aligned} \tag{3}$$

where now only the final term needs further simplification, since the denominator of the first part is independent of $c_i$, the numerator is the prior we have calculated previously and the middle term is independent of the cluster allocations.

We can rewrite this final term by introducing the following notation for sets of latent variables $z_j$:

$$\begin{aligned} C_l &:= \{z_j : j \neq i, c_j = l\} \\ \tilde{C}_l &:= \{z_j : c_j = l\} \end{aligned}$$

i.e. $\tilde{C}_l$ is the set of points assigned to cluster $l$ and $C_l$ is the same but excluding point $i$. We also introduce the notation that for a set $C$, $\langle C \rangle$ is the event that all the points in $C$ are generated from a single cluster. We can now rewrite the final term in Equation 3 as follows:

$$\begin{aligned} p(\boldsymbol{z}|c_i = k, \boldsymbol{c_{-i}}, \alpha, \theta_0) &= p(\left\langle \tilde{C}_1 \right\rangle, \ldots, \left\langle \tilde{C}_K \right\rangle |\alpha, \theta_0) \\ &= \prod_l p(\left\langle \tilde{C}_l \right\rangle |\alpha, \theta_0) \end{aligned}$$

where the second line follows by independence of clusters[1]. We can then split out the terms that depend on $k$ from those that will be the same in $p(\boldsymbol{z}|c_i = k, \boldsymbol{c_{-i}}, \alpha, \theta_0)$ for every $k$. Firstly, if $n_{-i,k} \neq 0$ then:

$$\begin{aligned} p(\boldsymbol{z}|c_i = k, \boldsymbol{c_{-i}}, \alpha, \theta_0) &= p(\langle C_k \cup \{z_i\} \rangle |\alpha, \theta_0) \prod_{l \neq k} p(\langle C_l \rangle |\alpha, \theta_0) \\ &= \frac{p(\langle C_k \cup \{z_i\} \rangle |\alpha, \theta_0)}{p(\langle C_k \rangle |\alpha, \theta_0)} \prod_{l=1}^{K} p(\langle C_l \rangle |\alpha, \theta_0) \end{aligned}$$

[1] Using the notation that $\langle C \rangle_\phi$ is the event that one of the clusters consists of the points in $C$ with parameter $\phi$:

$$\begin{aligned} p(\boldsymbol{z}|c_i = k, \boldsymbol{c_{-i}}, \alpha, \theta_0) &= \int_{\boldsymbol{\phi}} p(\left\langle \tilde{C}_1 \right\rangle, \ldots, \left\langle \tilde{C}_K \right\rangle |\boldsymbol{\phi}, \alpha, \theta_0) p(\boldsymbol{\phi}|\alpha, \theta_0) d\boldsymbol{\phi} \\ &= \int_{\boldsymbol{\phi}} p(\left\langle \tilde{C}_1 \right\rangle_{\phi_1}, \ldots, \left\langle \tilde{C}_K \right\rangle_{\phi_K} |\boldsymbol{\phi}, \alpha, \theta_0) \prod_j p(\phi_j|\alpha, \theta_0) d\boldsymbol{\phi} \\ &= \prod_j \int_{\phi_j} p(\left\langle \tilde{C}_j \right\rangle_{\phi_j} |\phi_j, \alpha, \theta_0) p(\phi_j|\alpha, \theta_0) d\phi_j \quad \text{By independence} \\ &= \prod_j p(\left\langle \tilde{C}_j \right\rangle |\alpha, \theta_0) \end{aligned}$$

and if $n_{-i,k} = 0$ i.e. if $k = K + 1$ so that this is a new cluster we have:

$$p(\boldsymbol{z}|c_i = k, \boldsymbol{c_{-i}}, \alpha, \theta_0) = p(\langle\{z_i\}\rangle\,|\alpha, \theta_0) \prod_{l=1}^{K} p(\langle C_l\rangle\,|\alpha, \theta_0)$$

In each case, the terms that do depend on $k$ are simply the marginal likelihoods of various sets of points ($C_k$, $C_k \cup \{z_i\}$ and $\{z_i\}$). Since we are using a conjugate prior for the cluster parameters $\phi = (\mu, \rho)$, these marginal likelihoods are easy to calculate (Section 3.3).

Thus, overall, we have the following equation for the conditional posterior of the cluster indicator variables:

$$p(c_i = k|\boldsymbol{c_{-i}}, \alpha, \boldsymbol{z}, \theta_0) = \begin{cases} \dfrac{p(\langle C_k \cup \{z_i\}\rangle\,|\alpha, \theta_0)}{p(\langle C_k\rangle\,|\alpha, \theta_0)} \dfrac{n_{-i,k}}{n - 1 + \alpha}\, b & \text{(if } n_{-i,k} \neq 0\text{)} \\ p(\langle\{z_i\}\rangle\,|\alpha, \theta_0) \dfrac{\alpha}{n - 1 + \alpha}\, b & \text{(if } n_{-i,k} = 0\text{)} \end{cases} \tag{4}$$

where $b$ is a constant that does not need to be calculated to sample $c_i$ from the conditional posterior to perform the Gibbs sampling step. For precision, we note that the constant $b$ is:

$$\frac{\prod_{l=1}^{K} p(\langle C_l\rangle\,|\alpha, \theta_0)}{p(\boldsymbol{z}, \boldsymbol{x}|\boldsymbol{c_{-i}}, \alpha, \theta_0)} p(\boldsymbol{x}|\boldsymbol{z})$$

#### 1.2 Distribution of cluster parameters

We perform Gibbs sampling on the cluster parameters for each cluster in turn, noting that cluster parameters are generated independently. We use the joint probability and discard terms that do not depend on the cluster parameters $\mu_k, \rho_k^2$. We show that that we simply have to sample from the posterior of a Normal-Inverse Gamma prior after 'observing' the latent data $z_{i_1}, \ldots, z_{i_{n_k}}$ in cluster $k$:

$$\begin{aligned} p(\mu_k, \rho_k^2|\boldsymbol{x}, \boldsymbol{z}, \boldsymbol{c}, \boldsymbol{\mu_{-k}}, \boldsymbol{\rho_{-k}}, \alpha) &= \frac{p(\boldsymbol{\mu}, \boldsymbol{\rho}, \boldsymbol{x}, \boldsymbol{z}, \boldsymbol{c}, \alpha)}{p(\boldsymbol{x}, \boldsymbol{z}, \boldsymbol{c}, \boldsymbol{\mu_{-k}}, \boldsymbol{\rho_{-k}}, \alpha)} \\ &= \frac{p(\boldsymbol{\mu}, \boldsymbol{\rho})p(\boldsymbol{c}|\alpha)p(\alpha)\prod_i p(x_i|\sigma_i^2, z_i)p(z_i|\mu_{c_i}, \rho_{c_i}^2)}{p(\boldsymbol{x}, \boldsymbol{z}, \boldsymbol{c}, \boldsymbol{\mu_{-k}}, \boldsymbol{\rho_{-k}}, \alpha)} \\ &\propto p(\mu_k, \rho_k^2) \prod_{i:c_i=k} p(z_i|c_i = k, \mu_k, \rho_k^2) \end{aligned}$$

noting that this last line is indeed the prior multiplied by the likelihood of the latent variables from this cluster, and is thus the posterior of a Normal-Gamma distribution which is itself a Normal-Gamma distribution.

#### 1.3 Distribution of latents

Again we can perform Gibbs sampling for each latent variable $z_i$ in turn. We again use the joint probability and discard the terms we do not need:

$$\begin{aligned} p(z_i|\boldsymbol{x}, \boldsymbol{z_{-i}}, \boldsymbol{c}, \boldsymbol{\mu}, \boldsymbol{\rho}, \alpha) &= \frac{p(\boldsymbol{x}, \boldsymbol{z}, \boldsymbol{c}, \boldsymbol{\mu}, \boldsymbol{\rho}, \alpha)}{p(\boldsymbol{x}, \boldsymbol{z_{-i}}, \boldsymbol{c}, \boldsymbol{\mu}, \boldsymbol{\rho}, \alpha)} \\ &= \frac{p(\boldsymbol{\mu}, \boldsymbol{\rho})p(\boldsymbol{c}|\alpha)p(\alpha)\prod_j p(x_j|\sigma_j^2, z_j)p(z_j|\mu_{c_j}, \rho_{c_j}^2)}{p(\boldsymbol{x}, \boldsymbol{z_{-i}}, \boldsymbol{c}, \boldsymbol{\mu}, \boldsymbol{\rho}, \alpha)} \\ &\propto p(x_i|\sigma_i^2, z_i)p(z_i|\mu_{c_i}, \rho_{c_i}^2) \end{aligned}$$

Thus we know that the conditional PDF of $z_i$ is proportional to the product of two Gaussian PDFs and so the conditional distribution is also a Gaussian distribution with parameters:

$$\sigma^2 = \frac{1}{\frac{1}{\sigma_i^2} + \frac{1}{\rho_k^2}}$$

and

$$\mu = \sigma^2 \left( \frac{z_i}{\sigma_i^2} + \frac{\mu_k}{\rho_k^2} \right)$$

### 2 Induced prior on $K$

DPMUnc does not directly place a prior on $K$, the number of clusters. However, there is a prior on $K$ induced by the rest of the model. Given that there are $n$ samples, and conditional on $\alpha$, we can write the expected number of clusters, $K$, as the following sum:

$$\mathbb{E}(K|\alpha, n) = \sum_{i=1}^{n} \frac{\alpha}{\alpha + i - 1}$$

since the $i$th individual has probability $\frac{\alpha}{\alpha+i-1}$ of forming a new cluster rather than joining an existing cluster.

We then make use of the digamma function, defined as the derivative of the natural logarithm of the gamma function:

$$\psi(x) = \frac{d}{dx} \ln(\Gamma(x)) = \frac{\Gamma'(x)}{\Gamma(x)} \sim \ln x - \frac{1}{2x}$$

For $x$ larger than 4 or so, even the $\frac{1}{2x}$ term is dwarfed by the ln term.

We use the property that $\psi(x+1) = \psi(x) + \frac{1}{x}$ as follows:

$$\begin{aligned}
\mathbb{E}(K|\alpha, n) &= \sum_{i=1}^{n} \frac{\alpha}{\alpha + i - 1} \\
&= \alpha \left[ \psi(\alpha) + \frac{1}{\alpha} + \frac{1}{\alpha+1} + \cdots + \frac{1}{\alpha+n-1} - \psi(\alpha) \right] \\
&= \alpha \left[ \psi(\alpha+1) + \frac{1}{\alpha+1} + \cdots + \frac{1}{\alpha+n-1} - \psi(\alpha) \right] \\
&\vdots \\
&= \alpha \left[ \psi(\alpha+n-1) - \psi(\alpha) \right] \\
&\sim \alpha \left[ \ln(\alpha+n-1) - \ln(\alpha) \right] \\
&\approx \alpha \ln\left( \frac{n}{\alpha} \right)
\end{aligned}$$

### 3 Further calculations for Gibbs equations

#### 3.1 Integrating out cluster weights

This section justifies Equation 1.

Recall that we have:

$$p(c_i = k) = \pi_k$$
$$\pi_1, \ldots, \pi_K \sim \text{Dirichlet}(\alpha/K, \ldots, \alpha/K)$$

We want to show that we can marginalise the cluster weights $\boldsymbol{\pi}$ from the prior on cluster indicators $c_i$. Thus instead of needing $p(\boldsymbol{c}|\alpha, \boldsymbol{\pi})$ we can simply use $p(\boldsymbol{c}|\alpha)$.

Recall that the PDF of a Dirichlet distribution with weights $\boldsymbol{\alpha} = (\alpha_1, \ldots, \alpha_K)$ is:

$$\frac{1}{\mathrm{B}(\boldsymbol{\alpha})} \prod_{k=1}^{K} \pi_k^{\alpha_k - 1}$$

where the normalising constant is:

$$\mathrm{B}(\boldsymbol{\alpha}) = \frac{\prod_{k=1}^{K} \Gamma(\alpha_k)}{\Gamma\left(\sum_{k=1}^{K} \alpha_k\right)}$$

Returning to our prior, we can integrate over $\boldsymbol{\pi}$ as follows:

$$\begin{aligned}
p(\boldsymbol{c}|\alpha) &= \int_{\boldsymbol{\pi}} p(\boldsymbol{c}|\alpha, \boldsymbol{\pi}) p(\boldsymbol{\pi}|\alpha) d\boldsymbol{\pi} \\
&= \int_{\boldsymbol{\pi}} \prod_{k=1}^{K} \pi_k^{n_k} \frac{\Gamma\left(\sum_{k=1}^{K} \frac{\alpha}{K}\right)}{\prod_{k=1}^{K} \Gamma\left(\frac{\alpha}{K}\right)} \prod_{k=1}^{K} \pi_k^{\frac{\alpha}{K} - 1} d\boldsymbol{\pi} \\
&= \frac{\Gamma(\alpha)}{\Gamma(\frac{\alpha}{K})^K} \int_{\boldsymbol{\pi}} \prod_{k=1}^{K} \pi_k^{n_k + \frac{\alpha}{K} - 1} d\boldsymbol{\pi}
\end{aligned}$$

We now recognise the integral as an unnormalised Dirichlet PDF with weights $\tilde{\boldsymbol{\alpha}} = (\tilde{\alpha}_1, \ldots, \tilde{\alpha}_K)$ where $\tilde{\alpha}_k = n_k + \frac{\alpha}{K}$ so we can use the usual trick of moving around the normalisation constants so that the integrand is a PDF and thus integrates to 1. Using the fact that the sum of these weights is $n + \alpha$ we can further simplify the equation as in the last step here:

$$\begin{aligned}
p(\boldsymbol{c}|\alpha) &= \frac{\Gamma(\alpha)}{\Gamma(\frac{\alpha}{K})^K} B(\tilde{\boldsymbol{\alpha}}) \cancel{\int_{\boldsymbol{\pi}} \frac{1}{B(\tilde{\boldsymbol{\alpha}})} \prod_{k=1}^{K} \pi_k^{\tilde{\alpha}_k - 1} d\boldsymbol{\pi}} \\
&= \frac{\Gamma(\alpha)}{\Gamma(\frac{\alpha}{K})^K} \frac{\prod_{k=1}^{K} \Gamma\left(n_k + \frac{\alpha}{K}\right)}{\Gamma(n + \alpha)}
\end{aligned}$$

##### 3.2 Prior of indicators conditional on other indicators

This section justifies Equation 2.

Using Equation 1 we can write the numerator as follows:

$$p(c_i = k, \boldsymbol{c_{-i}}|\alpha, K) = \frac{\Gamma(\alpha)}{\Gamma(n + \alpha)\Gamma\left(\frac{\alpha}{K}\right)^K} \Gamma\left(n_{-i,k} + 1 + \frac{\alpha}{K}\right) \prod_{l=1, l \neq k}^{K} \Gamma\left(n_{-i,l} + \frac{\alpha}{K}\right)$$

and then use the fact that $\Gamma(z+1) = z\Gamma(z)$ to make only one term dependent on $k$:

$$\begin{aligned}
p(c_i = k, \boldsymbol{c_{-i}}|\alpha, K) &= \frac{\Gamma(\alpha)}{\Gamma(n+\alpha)\Gamma\left(\frac{\alpha}{K}\right)^K}\Gamma\left(n_{-i,k}+\frac{\alpha}{K}\right)\left(n_{-i,k}+\frac{\alpha}{K}\right)\prod_{l=1,l\neq k}^{K}\Gamma\left(n_{-i,l}+\frac{\alpha}{K}\right) \\
&= \frac{\Gamma(\alpha)}{\Gamma(n+\alpha)\Gamma\left(\frac{\alpha}{K}\right)^K}\left(n_{-i,k}+\frac{\alpha}{K}\right)\prod_{l=1}^{K}\Gamma\left(n_{-i,l}+\frac{\alpha}{K}\right) \\
&= \left(n_{-i,k}+\frac{\alpha}{K}\right)C(\alpha, K, \boldsymbol{n_{-i}})
\end{aligned}$$

Every term in Equation 2 will have the constant $C(\alpha, K, \boldsymbol{n_{-i}})$ so we can cancel it and we are left with:

$$\begin{aligned}
p(c_i = k|\boldsymbol{c_{-i}}, \alpha, K) &= \frac{p(c_i = k, \boldsymbol{c_{-i}}|\alpha, K)}{\sum_{l=1}^{K} p(c_i = l, \boldsymbol{c_{-i}}|\alpha, K)} \\
&= \frac{n_{-i,k}+\frac{\alpha}{K}}{\sum_{l=1}^{K}\left(n_{-i,l}+\frac{\alpha}{K}\right)} \\
&= \frac{n_{-i,k}+\frac{\alpha}{K}}{n-1+\alpha}
\end{aligned}$$

##### 3.3 Marginal likelihood of latent data

We require the marginal likelihood $p(\langle C\rangle \mid \theta_0)$ for the Gibbs update for the cluster indicator variables (Equation 4), where we have grouped the constant hyperparameters for the cluster parameters into a single variable $\theta_0 = (\mu_0, \kappa_0, \alpha_0, \beta_0)$ for convenience, and group the cluster parameters into a single variable $\phi = (\mu, \rho)$.

Bayes' Theorem states:

$$p(\phi|\langle C\rangle, \theta_0) = \frac{p(\langle C\rangle|\phi, \theta_0)p(\phi|\theta_0)}{p(\langle C\rangle \mid \theta_0)}$$

We split each function into the part that depends on $\phi$, which we label $q$ and the constant part, which is the normalisation constant:

$$\frac{q(\phi|\langle C\rangle, \theta_0)}{Z_n} = \frac{q(\langle C\rangle|\phi, \theta_0)}{Z_l}\frac{q(\phi|\theta_0)}{Z_0}\frac{1}{p(\langle C\rangle \mid \theta_0)}$$

where $Z_n$ is the normalisation constant for the posterior, $Z_0$ is the normalisation constant for the prior and $Z_l$ is the normalisation constant for the likelihood.

Thus by equating the constant terms we get a simple equation for the marginal likelihood:

$$p(\langle C\rangle \mid \theta_0) = \frac{Z_n}{Z_0 Z_l}$$

The latent data have Gaussian distribution, with prior for $\mu_k$ and $\tau = 1/\rho_k^2$ having Normal-Gamma distribution. If a prior Normal-Gamma distribution has parameters $\mu_0, \kappa_0, \alpha_0, \beta_0$ then the posterior after seeing observations $z_1, \ldots, z_n$ will also be a Normal-Gamma distribution, with parameters $(\mu_n, \kappa_n, \alpha_n, \beta_n)$ given by:

$$\begin{aligned}
\mu_n &= \frac{\kappa_0 \mu_0 + n\bar{z}}{\kappa_0 + n} \\
\kappa_n &= \kappa_0 + n \\
\alpha_n &= \alpha_0 + \frac{n}{2} \\
\beta_n &= \beta_0 + \frac{1}{2}\left(\text{RSS} + \frac{\kappa_0 n \left(\bar{z} - \mu_0\right)^2}{\kappa_0 + n}\right)
\end{aligned}$$

where $\bar{z} = \frac{1}{n}\sum_i z_i$ is the empirical mean and $\text{RSS} = \sum_i (z_i - \bar{z})^2$ is the residual sum of squares.

So $Z_0$ is simply the normalisation constant of a Normal-Gamma distribution with parameters $(\mu_0, \kappa_0, \alpha_0, \beta_0)$, and $Z_n$ is simply the normalisation constant of a Normal-Gamma with parameters $(\mu_n, \kappa_n, \alpha_n, \beta_n)$. Since the latent data are i.i.d. with Gaussian distribution, the normalisation constant of the likelihood is just the normalisation constant of a Gaussian, which is $Z_l = (2\pi)^{\frac{-n}{2}}$.

Thus overall, a set of latent variables $C$ has marginal likelihood:

$$p(\langle C \rangle \mid \theta_0) = \frac{\beta_n^{\alpha_n}\sqrt{\kappa_n}}{\Gamma(\alpha_n)\sqrt{2\pi}} \frac{\Gamma(\alpha_0)\sqrt{2\pi}}{\beta_0^{\alpha_0}\sqrt{\kappa_0}} \frac{1}{(2\pi)^{\frac{-n}{2}}}$$

where $n$ is the number of elements in $C$.

Note that since the location parameter $\mu$ does not appear in the normalisation constant of a Normal-Gamma distribution, we do not actually need to calculate $\mu_n$.

#### 4 Choice of $\alpha_0, \beta_0$

The conjugate prior for a Gaussian with unknown mean and variance is a Normal-Gamma. Our model can thus be written as follows:

$$\begin{aligned} z|\mu,\tau &\sim \mathcal{N}(\mu,\tau^{-1}) \\ \mu|\tau &\sim \mathcal{N}(\mu_0,(\kappa_0\tau)^{-1}) \\ \tau|\alpha_0,\beta_0 &\sim \text{Gamma}(\alpha_0,\beta_0) \end{aligned} \tag{5}$$

using $\tau = \frac{1}{\rho^2}$.

After some experimentation, we found that if the variance of $z$ is fixed to be 1, a relatively general prior uses $\alpha_0 = 2, \beta_0 = 0.2, \kappa_0 = 0.1$.

We can now adjust this to data which has not been scaled to have variance 1. We want $\tilde{z} := z/s$, where $s$ is the standard deviation of $z$, to have the distribution given in Equation 5.

$$\begin{aligned} \frac{z}{s} \mid \mu,\tau \sim \mathcal{N}(\mu,\tau^{-1}) &\implies z|\mu,\tau \sim \mathcal{N}(\mu s, s^2\tau^{-1}) \\ &\implies z|\mu,\tau \sim \mathcal{N}(\mu s, (\frac{\tau}{s^2})^{-1}) \end{aligned} \tag{6}$$

We can rewrite this as:

$$z|\tilde{\mu},\tilde{\tau} \sim \mathcal{N}(\tilde{\mu},\tilde{\tau}^{-1})$$

where $\tilde{\mu} = \mu s$ and $\tilde{\tau} = \frac{\tau}{s^2}$ and we can continue to rewrite the equations from Equation 5:

$$\begin{aligned} \mu|\tau \sim \mathcal{N}(\mu_0,(\kappa_0\tau)^{-1}) &\implies s\mu|\tau \sim \mathcal{N}(s\mu_0, (\frac{\kappa_0\tau}{s^2})^{-1}) \\ &\implies \tilde{\mu}|\tilde{\tau} \sim \mathcal{N}(\tilde{\mu_0},(\tilde{\kappa_0}\tilde{\tau})^{-1}) \end{aligned} \tag{7}$$

with $\tilde{\mu_0} = s\mu_0$ and $\tilde{\kappa_0} = \kappa_0$.

Similarly with the final equation

$$\begin{aligned} \tau|\alpha_0,\beta_0 \sim \mathcal{N}(\alpha_0,\beta_0) &\implies \frac{\tau}{s^2}|\alpha_0,\beta_0 \sim \text{Gamma}(\alpha_0, s^2\beta_0) \\ &\implies \tilde{\tau}|\tilde{\alpha}_0,\tilde{\beta}_0 \sim \text{Gamma}(\tilde{\alpha}_0,\tilde{\beta}_0) \end{aligned} \tag{8}$$

where $\tilde{\alpha}_0 = \alpha_0$ and $\tilde{\beta}_0 = s^2\beta_0$. So overall the model is:

$$\begin{aligned} z|\tilde{\mu},\tilde{\tau} &\sim \mathcal{N}(\tilde{\mu},\tilde{\tau}^{-1}) \\ \tilde{\mu}|\tilde{\tau} &\sim \mathcal{N}(\tilde{\mu}_0,(\tilde{\kappa_0}\tilde{\tau})^{-1}) \\ \tilde{\tau}|\tilde{\alpha}_0,\tilde{\beta}_0 &\sim \text{Gamma}(\tilde{\alpha}_0,\tilde{\beta}_0) \end{aligned} \tag{9}$$

Thus after reparameterisation we find that the only change we need to make is to multiply $\beta_0$ by the variance of $z$, which is $s^2$, and to multiply $\mu_0$ by $s$. In practice, we set $\mu_0$ to the empirical mean of $x$, which is $\bar{x}$.

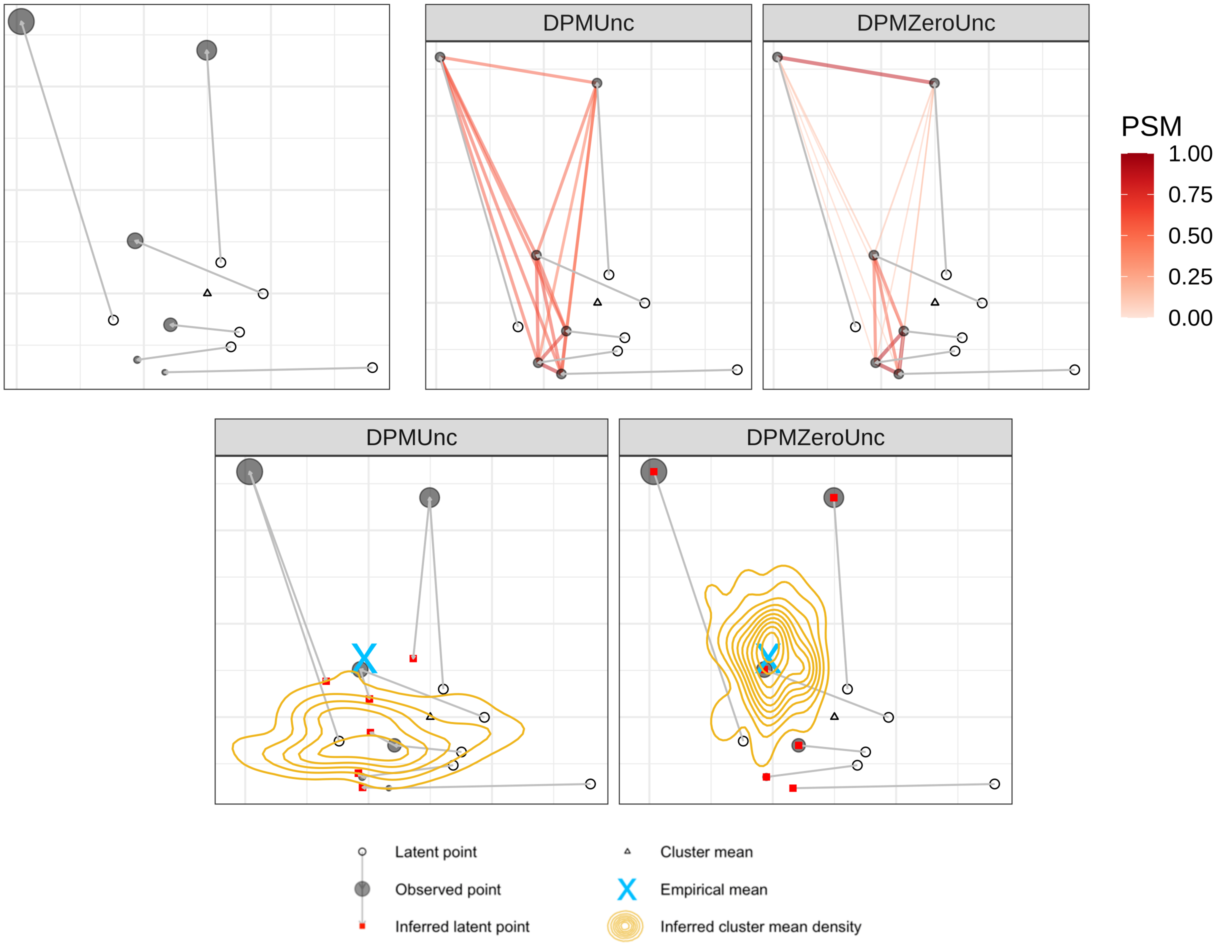


Supplementary Figure S1: Top left: example *mean shift* dataset to illustrate how the inferred cluster mean can shift when uncertainty is taken into account. The hollow circles represent true underlying latent points, which would be unknown for a real dataset. An arrow connects each point to the corresponding observed value, whose size represents the amount of uncertainty associated with the observation. The triangle is the true cluster mean. Top middle and right: posterior similarity scores. Note that the final output of both DPMUnc and DPMZeroUnc was a single cluster containing all the points. Each pair of points is connected by a line whose width and darkness grows with the posterior similarity score between the points: the proportion of samples taken by DPMUnc in which the two points were clustered together. Bottom row: posterior distribution of cluster mean, when the clustering contains a single cluster. A further arrow connects the observed point to a red square which is the mean latent observation inferred by DPMUnc (or DPMZeroUnc). The blue 'X' is the empirical mean of the observations and the golden densities show the distribution of the cluster mean inferred by each method.

### 5 Illustrative examples

#### 5.1 Shift in cluster mean

The *mean shift* example contains six observations in a single cluster, with two observations lying far from the corresponding latent points and from the cluster mean due to high observational noise (Figure S1, top row). Without taking the uncertainty of the points into account, these outlying points are at risk of being placed in a separate cluster. Indeed, the k-means solution for this dataset is to place these two points in one cluster, and the remaining points in another cluster. The mclust solution has five clusters. The clustering returned by DPMZeroUnc does in fact put all points in the same cluster, but the posterior similarity scores are much lower between each of these two points and the remaining points, suggesting that in some samples from DPMZeroUnc, the two points with high observational noise were placed in a separate cluster. It is instructive to look at the latent variables that DPMUnc inferred and the distribution of the cluster mean when DPMUnc inferred a single cluster (Figure S1, bottom row). DPMUnc inferred latent variables closer to the empirical mean of the points that had low uncertainty. In samples from DPMUnc consisting of a single cluster, the inferred cluster mean was also shifted closer to these points with lower uncertainty, whereas when DPMZeroUnc inferred a single cluster, the cluster mean was very close to the empirical mean of all the observed points. Note also that the distribution of the cluster mean when uncertainty is taken into account is much less concentrated on a single value, because the uncertainty of the exact locations of the data leads to uncertainty in DPMUnc's sampling.

Thus the *mean shift* example dataset illustrates that DPMUnc can infer latent points very different from the observed points, and that when there is a range of uncertainty in the dataset the cluster mean gets shifted towards the points that have lower uncertainty. It also illustrates that confidence in the exact location of the cluster mean is also (justifiably) reduced when uncertainty is taken into account.

#### 5.2 Shrinking of cluster variance

The *cluster shrinkage* example dataset consists of 40 points across two clusters, generated using the simulation procedure described in Methods, with high uncertainty about each observation $U = 10$, low cluster noise $N = 2$ and cluster means at $(20, 0)$ and $(0, 20)$ (Figure S2). The clustering itself is very simple and all four methods recovered the true clustering exactly. However, DPMUnc additionally estimates the underlying latent variables and these estimates provide an insight into the difference made by taking uncertainty into account. The estimates of the latent variables provided by DPMUnc have significantly lower variance than the observed data (Figure S2). Further, the median cluster variance inferred by DPMUnc (for the samples where DPMUnc inferred two clusters) was $(12.188, 10.695)$ whereas DPMZeroUnc had median cluster variance $(12.997, 11.044)$. The full distribution of inferred cluster variances for DPMUnc and DPMZeroUnc are shown in Figure S5.

Thus in this *cluster shrinkage* example dataset we observe that the latent variables move towards the cluster center and the estimated cluster variance is smaller than when uncertainty is not taken into account. The clusters in this illustrative example are so well separated that the clustering itself is not impacted. However, in more complicated datasets, the movement of the latent variables, and the shrinkage of the cluster variance could change the clustering recovered by DPMUnc.

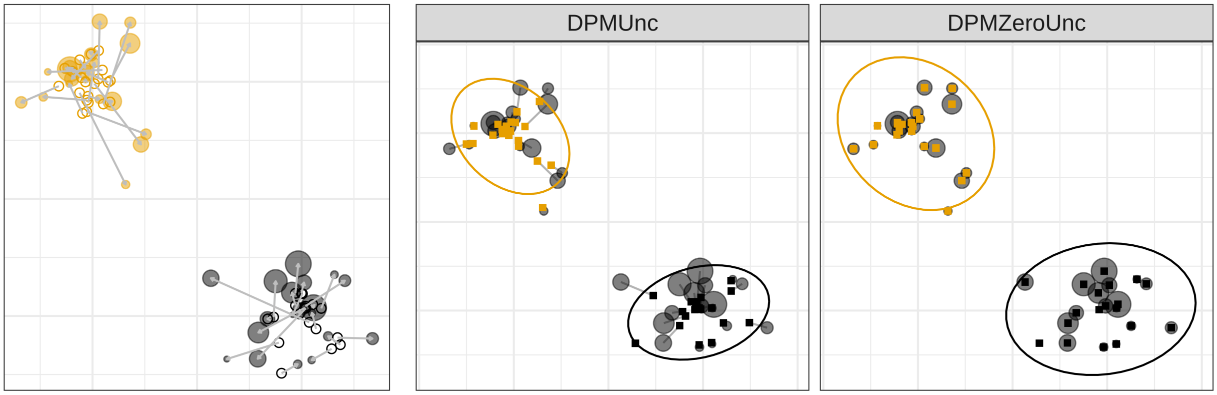


Supplementary Figure S2: The left panel shows the *cluster shrinkage* dataset. The hollow circles represent true underlying latent points, which would be unknown for a real dataset. The grey circles are the observed values, whose size represents the amount of uncertainty associated with the observation. An arrow connects each latent point to the corresponding observed value. The next panels show the cluster solutions, allowing for uncertainty (DPMUnc, middle) and ignoring it (DPZeroMunc, right). An arrow connects each observation to the latent variable inferred by DPMUnc or DPMZeroUnc, represented by squares, noting that DPMZeroUnc infers latent variables equal to the observations since it does not take uncertainty into account. The colour of the inferred latents corresponds to the true cluster. Each cluster is surrounded by an ellipse indicating the 95% confidence interval for the cluster, i.e. the ellipse is approximately mean $\pm$ 1.96 sd.

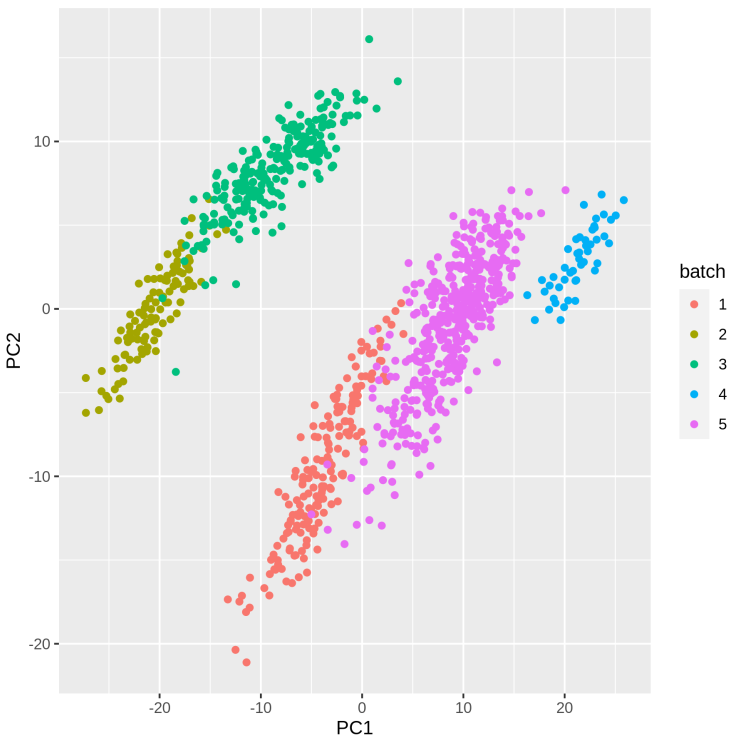


Supplementary Figure S3: Principal component analysis of the samples in E-MTAB-1724, showing a clear batch effect. We take batch 1 to be the Ferreira dataset.

#### References

| $Sig_{\text{IFN}}$ | | | 56 genes | |
|---|---|---|---|---|
| ANKRD22 | BRCA2 | CMPK2 | CXCL10 | CXCL11 |
| DDX58 | DDX60 | DHX58 | DTX3L | EIF2AK2 |
| EPSTI1 | ETV7 | FAM70A | HERC5 | HERC6 |
| HMCN2 | IFI35 | IFI44 | IFI44L | IFIH1 |
| IFIT1 | IFIT2 | IFIT3 | IFIT5 | IRF7 |
| ISG15 | ISG20 | LAMP3 | LAP3 | LGALS3BP |
| LY6E | MX1 | MX2 | NEXN | OAS1 |
| OAS2 | OAS3 | OASL | OTOF | PARP12 |
| PARP9 | PGAP1 | PML | PNPT1 | RGL1 |
| RP11-445H22.3 | RP4-697K14.7 | RSAD2 | SAMD9L | SIGLEC1 |
| SPATS2L | STAT1 | STAT2 | USP18 | USP41 |
| XAF1 | | | | |
| $Sig_{\text{NK}}$ | | | 87 genes | |
| KLRG1 | NMUR1 | GNLY | CXCR6 | YME1L1 |
| PRSS23 | CD160 | SH2D1B | CLDND2 | PYHIN1 |
| CTSW | ADRB2 | SAMD3 | APOBEC3H | TOGARAM2 |
| DUSP2 | PATL2 | TIGIT | ERBB2 | SYT11 |
| MCOLN2 | GK5 | ZNF683 | NCR3 | GFI1 |
| STK39 | GNGT2 | TSEN54 | YPEL1 | GZMH |
| GZMB | ST6GALNAC6 | FCRL6 | FASLG | IL2RB |
| KIR2DL3 | KLRC1 | KLRD1 | LAG3 | MAF |
| MATK | TARP | MYBL1 | NCAM1 | NKG7 |

| | | | | |
|---|---|---|---|---|
| RNF165 | KLRF1 | PDGFRB | S1PR5 | TTC38 |
| PPP2R2B | CHST12 | PRF1 | KIF21A | LYAR |
| PTGDR | PDZD4 | PTPN4 | SLAMF7 | ABHD17C |
| EVA1C | LGR6 | CCL5 | EPB41L4A | DNAJC1 |
| SLC20A1 | STAT4 | TGFBR3 | PLEKHF1 | PRR5L |
| PDGFD | GPR68 | EOMES | FGFBP2 | NCALD |
| HOPX | TPST2 | RUNX3 | CLIC3 | SH2D2A |
| LDB2 | CD247 | TSPOAP1 | ADGRG1 | SOX13 |
| FEZ1 | JAKMIP2 | | | |
| $Sig_{\mathrm{CD8}}$ | | | 12 genes | |
| STK38 | ITGB7 | KLF3 | DGKA | EMB |
| EPHB4 | C5orf34 | SDHA | SATB1 | NFE2L2 |
| ADH5 | EEF2 | | | |

Table S1: Genes in each signature used in the gene expression datasets.

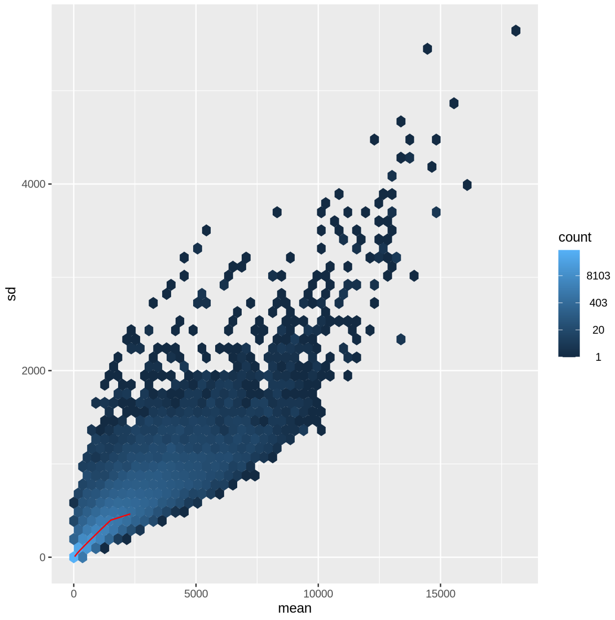


(a) Raw data

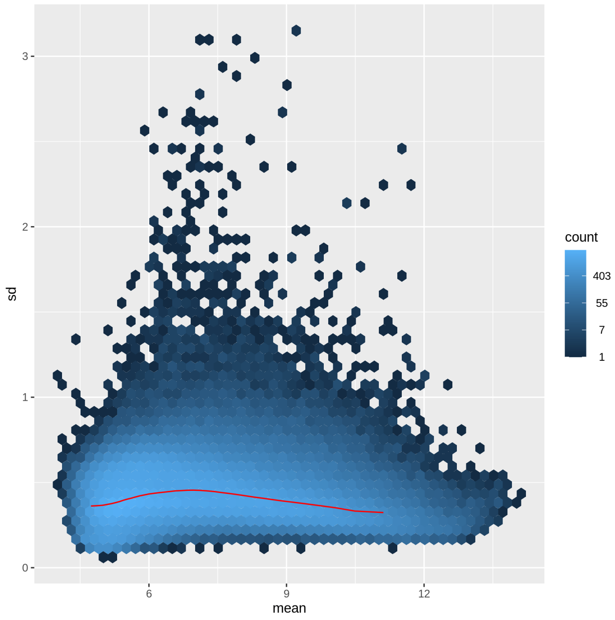


(b) After vsn2

Supplementary Figure S4: Relationship between mean and standard deviation in the Ferreira dataset, before and after vsn2 transformation.

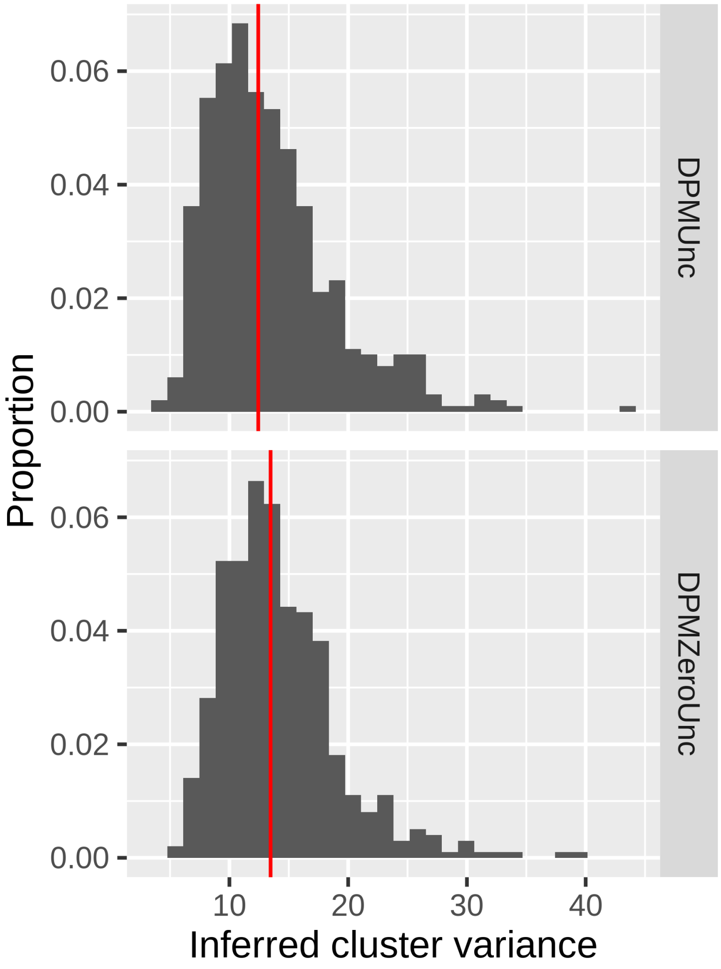


Supplementary Figure S5: Distribution of inferred cluster variances on the *cluster shrinkage* dataset. The median for each method is shown with a red line.

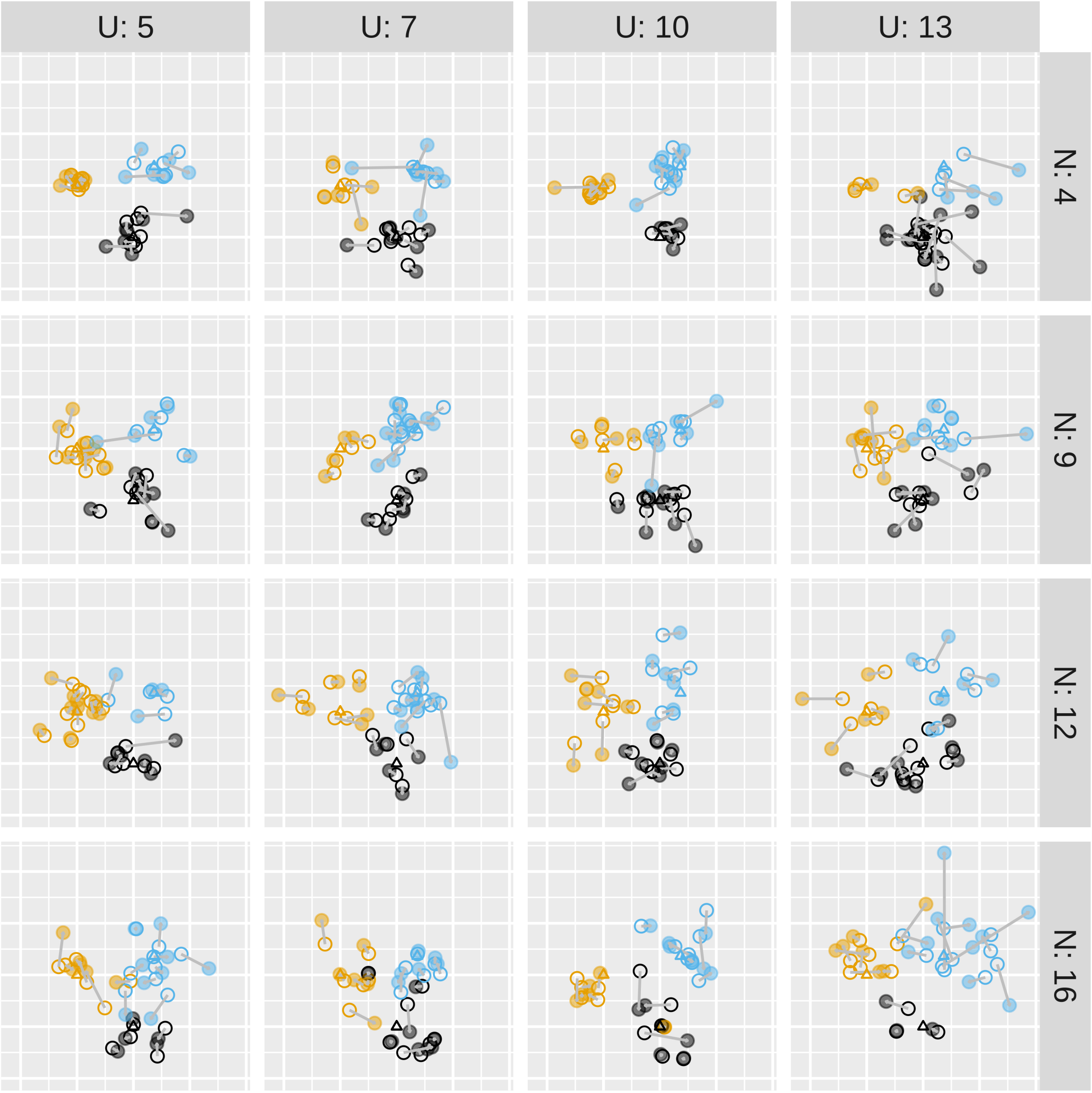


Supplementary Figure S6: Example simulated datasets with varying levels of true noise $N$ and observation uncertainty $U$. The latent data are empty circles, with arrows pointing to the observed values, which are represented by filled circles. The points are coloured by cluster membership. The left column has lowest observation uncertainty, so that the observations are close to their latent points, whereas the right column has high observation uncertainty. The top row has lowest cluster noise, with most latent data lying close to the cluster mean whereas the bottom row has higher cluster noise, with latent data more spread out.

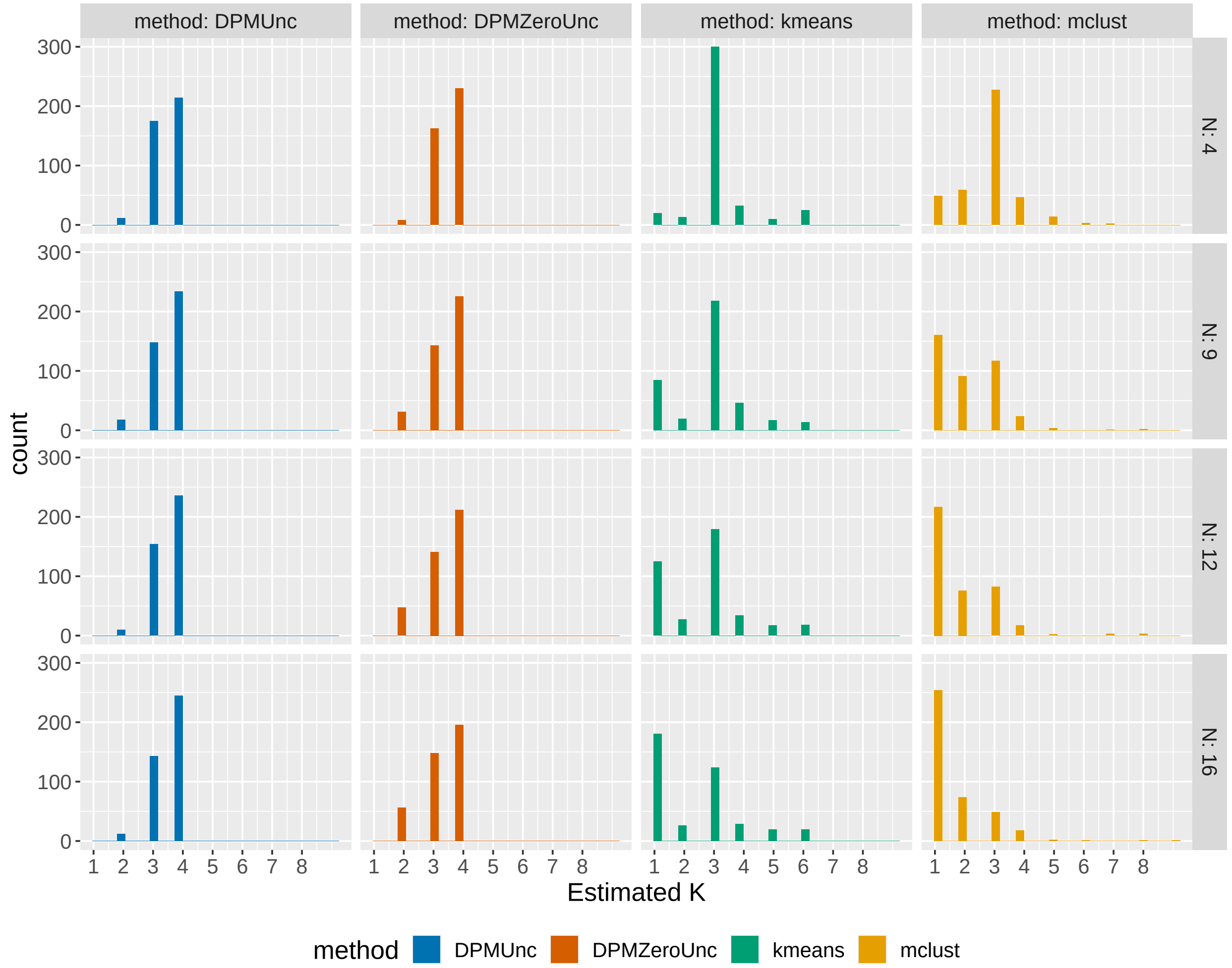


Supplementary Figure S7: Estimates of $K$ for different methods across simulated datasets. In all cases true K is 3.

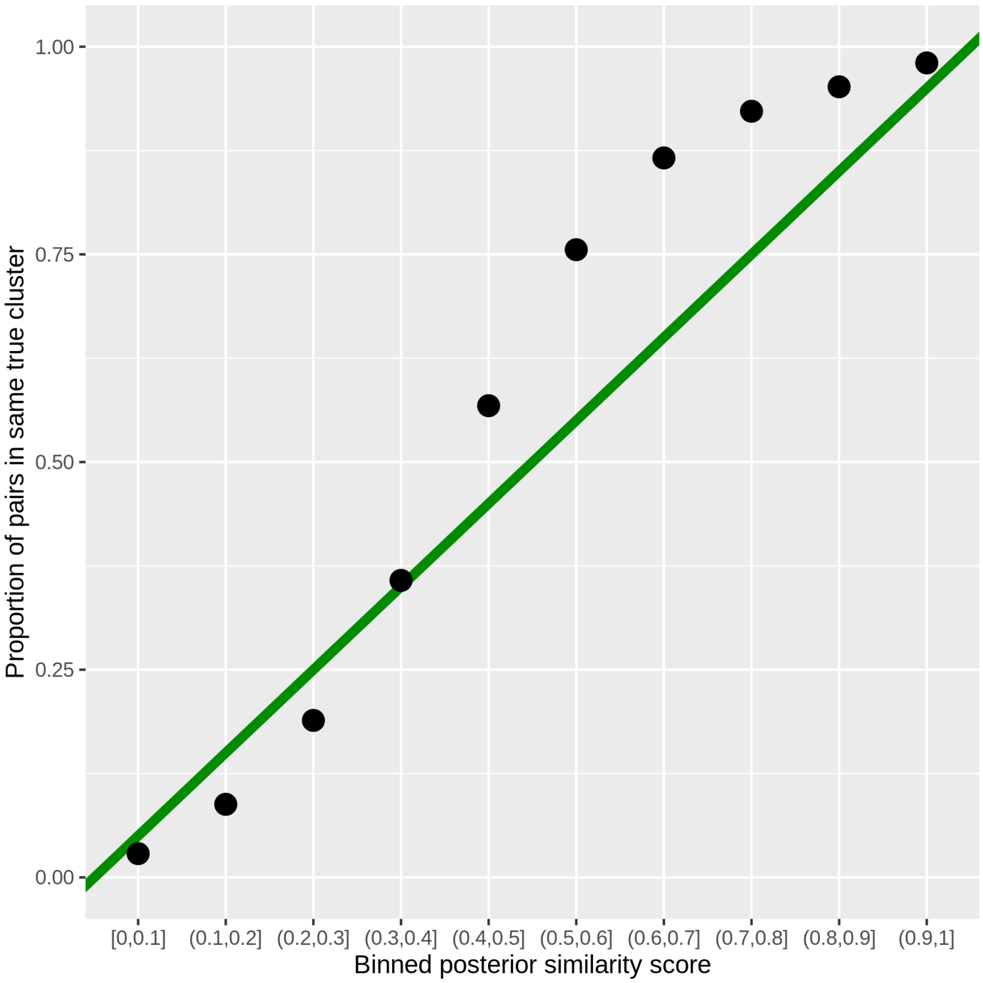


Supplementary Figure S8: Calibration of posterior similarity scores. The green line shows perfect calibration, where the posterior similarity score is the same as the proportion of pairs that are in the same cluster in the true clustering. For each collection of pairs of points with posterior similarity score within some bin, the proportion of pairs which are in the same cluster in the true clustering is shown by a dot.

Supplementary Figure S9: MCMC algorithm used by DPMUnc

$z \leftarrow x$
**for** $iter = 1, \ldots, numIterations$ **do**
  **for** $z_i : i = 1, \ldots n$ **do**
    **for** $k = 1, \ldots K$ **do**
      $w_k = \frac{|C_k \setminus z_i|}{n-1+\alpha} \frac{p(C_k \cup z_i)}{p(C_k \setminus z_i)}$ ▷ Ratio of marginal likelihood of cluster with and without the given observation
    **end for**
    $w_{K+1} = \frac{\alpha}{n_k+\alpha} p(z_i)$
    $p(c_i = k) = w_k$
  **end for**
  $\alpha \sim p(\alpha|K)$
  $\mu_k, \rho_k^2 \sim p(\mu_k, \rho_k^2 \mid \{x_i : z_i = k\})$ ▷ Posterior of cluster variables is Normal-Inverse Gamma
  $z_i \overset{\text{i.i.d.}}{\sim} p(z_i \mid \mu_k, \rho_k^2, x, z_i = k)$ ▷ Posterior of latent data is Normal
**end for**

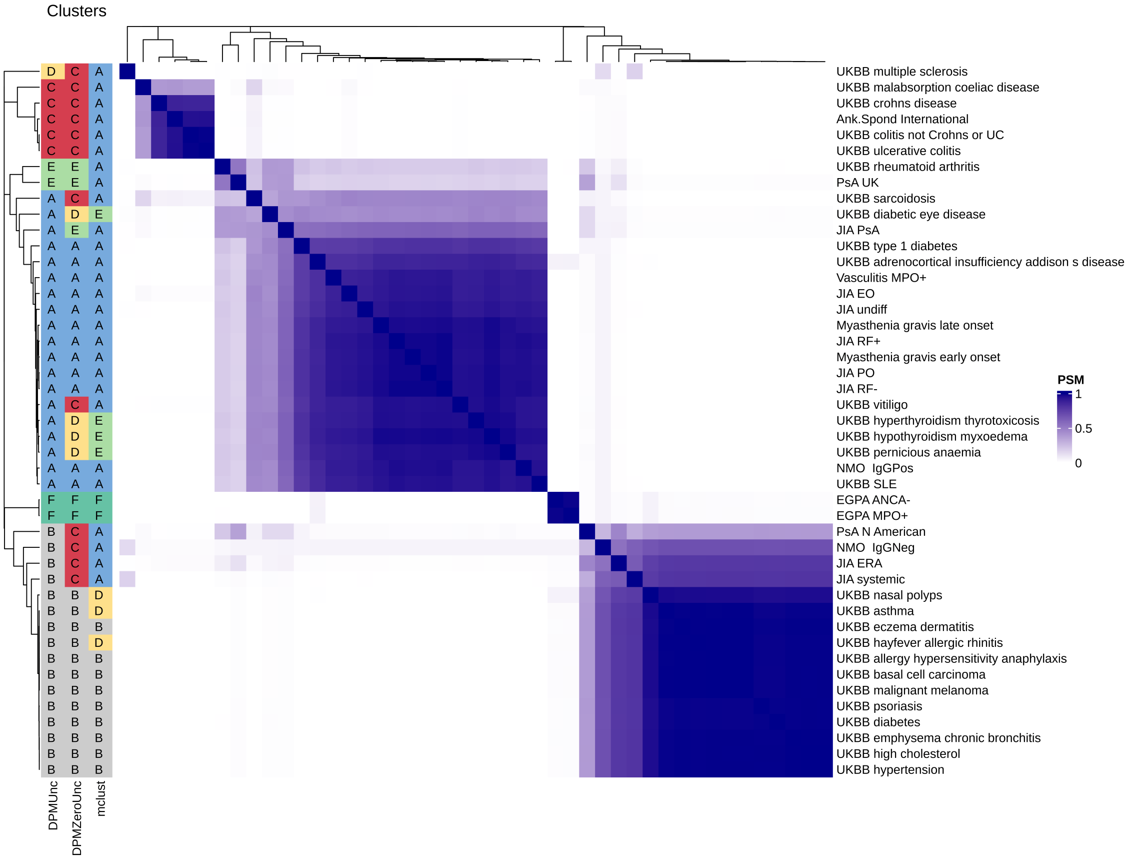


Supplementary Figure S10: Posterior similarity matrix for DPMUnc's clustering. Both rows and columns correspond to traits and each entry in the grid shows the proportion of samples from the MCMC chain in which the two traits were placed in the same cluster.
